# Accurate Reconstruction of Microbial Strains from Metagenomic Sequencing Using Representative Reference Genomes

**DOI:** 10.1101/215707

**Authors:** Zhemin Zhou, Nina Luhmann, Nabil-Fareed Alikhan, Christopher Quince, Mark Achtman

## Abstract

Exploring the genetic diversity of microbes within the environment through metagenomic sequencing first requires classifying these reads into taxonomic groups. Current methods compare these sequencing data with existing biased and limited reference databases. Several recent evaluation studies demonstrate that current methods either lack sufficient sensitivity for species-level assignments or suffer from false positives, overestimating the number of species in the metagenome. Both are especially problematic for the identification of low-abundance microbial species, e. g. detecting pathogens in ancient metagenomic samples. We present a new method, SPARSE, which improves taxonomic assignments of metagenomic reads. SPARSE balances existing biased reference databases by grouping reference genomes into similarity-based hierarchical clusters, implemented as an efficient incremental data structure. SPARSE assigns reads to these clusters using a probabilistic model, which specifically penalizes non-specific mappings of reads from unknown sources and hence reduces false-positive assignments. Our evaluation on simulated datasets from two recent evaluation studies demonstrated the improved precision of SPARSE in comparison to other methods for species-level classification. In a third simulation, our method successfully differentiated multiple co-existing *Escherichia coli* strains from the same sample. In real archaeological datasets, SPARSE identified ancient pathogens with *≤* 0.02% abundance, consistent with published findings that required additional sequencing data. In these datasets, other methods either missed targeted pathogens or reported non-existent ones. SPARSE and all evaluation scripts are available at https://github.com/zheminzhou/SPARSE.

## 1 Introduction

Shotgun metagenomics generates DNA sequences directly from environmental samples, revealing unculturable organisms in the community as well as those that can be isolated. The resulting data represents a pool of all species within a sample, thus raising the problem of identifying individual microbial species and their relative abundance within these samples. Methods for such taxonomic assignment are either based on *de novo* assembly of the metagenomic reads, or take advantage of comparisons to existing *reference* genomes. Here we concentrate on the latter strategy, which relies on the diversity of genomes in ever-growing reference databases. This strategy has been instrumental in identifying many causative agents of ancient pandemics in reads obtained from archaeological samples by detecting genetic signatures of modern human pathogens [26].

Published methods for taxonomic assignment can be divided into two categories. *Taxonomic profilers* maintain a small set of curated genomic markers, which can be universal (e. g. used in MIDAS [16]) or clade-specific (e. g. used in MetaPhlan2 [24]). Metagenomic reads that align onto these genomic markers are used to extrapolate the taxonomic composition of the whole sample. These tools are usually computationally efficient with good precision. However, they also tend to show reduced resolution for species-level assignment [23], especially when a species has a low abundance in the sample and, hence, may have few reads mapping to a restricted set of markers.

Alternatively, *taxonomic binners* compare metagenomic reads against reference genomes to achieve read-level taxonomic classification. The comparisons can be kmer-based (e. g. Kraken [25] and One Codex [15]) or alignment-based (MEGAN [6], MALT [5] and Sigma [1]). Binning methods based on kmers are usually fast, whilst alignment-based methods have greater sensitivity to distinguish the best match across similar database sequences. Benefiting from much larger databases in comparison to genomic markers used by profiling methods, binning methods usually detect more microbial species at very low abundance. However, they also tend to accumulate inaccurate assignments (false positives) [23] due to the incompleteness of the databases, resulting in reads from unrepresented taxa being erroneously attributed to multiple relatives.

While microbial species of low abundance are hard to identify by marker-based taxonomic profilers, the estimations of taxonomic binners can be hard to interpret due to their low precision. This problem especially limits their application to the *in silico* screening of microbial content in sequenced archaeological materials [8]. Given that the ancient DNA fragments are expected to exist in low proportions in these samples, methods need to identify weak endogenous signatures hidden within a complex background that is governed by modern (environmental) contamination. Furthermore, reads from archaeological samples are fragmented and have many nucleotide mis-incorporations due to postmortem DNA damage.

We identify two challenges that limit the performance of species-level assignments. First and foremost, the reference database used for all taxonomic binnings are not comprehensive. The vast majority of microbial genetic diversity reflect uncultured organisms, which have only rarely been sequenced and analyzed. Even for the bacteria that have genomic sequences, their data are biased towards pathogens over environmental species. This leads to the next challenge where, due to the lack of proper references, reads from unknown sources can accidentally map onto distantly related references, mainly in two scenarios: 1) Foreign reads originating from a mobile element can non-specifically map to an identical or similar mobile element in a known reference. 2) Reads originated from Ultra-Conserved Elements (UCEs), which preserve their nucleotide sequences between species, can also non-specifically map to the same UCE in an existing genome.

Addressing both of these challenges, we designed SPARSE (**S**train **P**rediction and **A**nalysis using **R**epresentative **SE**quences). In SPARSE, we index all genomes in large reference databases such as RefSeq into hierarchical clusters based on different sequence identity thresholds. A representative database that chooses one sequence for each cluster is then compiled to facilitate a fast but sensitive analysis of metagenomic samples with modest computational resources. Details are given in Section 2. Further, SPARSE implements a probabilistic model for sampling reads from a metagenomic sample, which extends the model described in Sigma [1] by weighting each read with its probability to stem from a genome not included in the reference database, hence considered as an unknown source. Details are given in Section 3.

We evaluate SPARSE on three simulated datasets published previously [14, 21, 23]. Comparing SPARSE to several other taxonomic binning software in these simulations shows its improved precision and sensitivity for assignments on the species-level or even strain-level. We further evaluate SPARSE on three ancient metagenomic datasets, demonstrating the application of SPARSE for ancient pathogen screening. For all three datasets, SPARSE is able to correctly identify small amounts of ancient pathogens in the metagenomic samples that have subsequently been confirmed by additional sequencing in the respective studies.

## 2 Database indexing

### 2.1 Background

#### Average nucleotide identity

To catalog strain-level genomic variations within an evolutionary context, we need to reconcile all the references in a database into comprehensive classifications. Since its first publication, the average nucleotide identity (ANI) in the conserved regions of genomes has been widely used for such a purpose [10]. In particular, 95 *−* 96% ANI roughly corresponds to a 70% DNA-DNA hybridization value, which has been used for ~ 50 years as the definition for prokaryotic species.

Marakeby et al. [13] proposed a hierarchical clustering of individual genomes based on multiple levels of ANIs. Extending from the 95% ANI species cut-off, it allows the classification of further taxonomic levels from superkingdoms to clones. However, the standard ANI computation adopts BLASTn [2] to align conserved regions between genomes, which is intractable to catalog large databases of reference genomes. We therefore rely on an approximation of the ANI by MASH [18] to speed-up comparisons.

#### ANI approximation

MASH uses the MinHash dimensionality-reduction technique to reduce large genomes into compressed sketches. A sketch is based on a hash function applied to a kmer representation of a genome, and compression is achieved by only including the *s* smallest hash values of all kmers in the genome in the sketch. Comparing the sketches of two genomes, MASH defines a distance measure under a simple Poisson process of random site mutation that approximates ANI values as shown in [18].

#### Parameter estimation

Ondov et al. [18] already used MASH to group all genomes in RefSeq into ANI 95% clusters. We adopted slightly different parameters and extended it to an incremental, hierarchical clustering system. The accuracy of the MASH distance approximation is determined by both the kmer length *k* and the sketch size *s*. Increasing *k* can reduce the random collisions in the comparison but also increase the uncertainty of the approximation. We can determine *k* according to equation (2) in [18]:

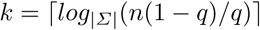

where *Σ* is the set of all four possible nucleotides {*A, C, G, T*}, *n* is the total number of nucleotides and *q* is the allowed probability of a random kmer to be found in a dataset. Given *n* = 1 terabase-pairs (Tbp; current size of RefSeq) and *q* = 0.05, which allows a 5% chance for a random k-mer to be present in a 1 Tbp database, we obtain a desired kmer size *k* = 23. Increasing the sketch size *s* will improve the accuracy of the approximation, but will also increase the run time linearly. We chose *s* = 4000 such that for 99.9% of comparisons that have a MASH distance of 0.05, the actual ANI values fall between 94.5 *−* 95.5%.

### 2.2 SPARSE reference database

We combine the hierarchical clustering of several ANI levels with the MASH distance computation to generate a representation of the current RefSeq [17] database. The construction of the SPARSE reference database is parallelized and incremental, thus the database can be easily updated with new genomes without a complete reconstruction.

#### Hierarchical clustering

In order to cluster genomes in different levels, we defined 8 different ANI values *L* = [0.9, 0.95, 0.98, 0.99, 0.995, 0.998, 0.999, 0.9995], in which the genetic distances of two sequential levels differ by ~ 2 fold. The first four ANI levels differentiate strains of different species, or major populations within a species. The latter four levels give fine-grained resolutions for intra-species genetic diversities, which can be used to construct clade-specific databases for specific bacteria.

The SPARSE database *D*(*S, L, K*) is extended incrementally as shown in Algorithm 1, with *S* listing the sketches of all genomes already in the database and *K* being a hash containing the cluster assignments at each level *l ∈ L* for each key *s ∈ S*. A new genome is integrated by finding another genome in the database with the lowest distance using MASH, and clustering it with its nearest neighbour *s*_*n*_ depending on the ANI.

##### Algorithm 1 Incremental SPARSE database clustering

**Input:** SPARSE database *D*(*S, L, K*), list of new genomes *G*

**Output:** Extended SPARSE database *D*^′^(*S, L, K*)

1. **for each** genome *g ∈ G* **do**
2. *s*_*g*_ = MashSketch(g)
3. *s*_*n*_ = *argmin*_*s∈S*_*MashDistance(s*_*g*_, *s*)
4. **for** 0 ≤ *i* ≤ |*L*| - 1] **do**
5. **if** *L[i]* ≤ *1 - MashDistance(s*_*g*_,*s*_*n*_) **then**
6. Push *K*[*s*_*n*_]*[i]* to *K*[*s*_*g*_]
7. **else**
8. Push |*S*| to *K[s*_*g*_]
9. Push *s*_*g*_ to *S*

In the SPARSE implementation, we parallelized the database construction by inserting batches of genomes at once and parallelizing sketch and distance computation, thereby scaling to the complexity of the problem. After being added to the database, the cluster assignment for a genome is fixed and never redefined. Therefore, the insertion order of genomes can influence the database structure. Here we utilize prior knowledge from the community, so the SPARSE database is initialized first with all *gold standard* complete genomes in RefSeq, followed by representative and curated genomes. With this strategy, the whole RefSeq database with 101, 680 genomes (Aug. 2017) can be downloaded and assigned into ANI levels in ~ 23hrs, using 20 processes on a standalone server. Further insertion of 1, 000 new genomes (~ 5MB) into an already established database takes ~ 15mins.

#### Representative database

To avoid mapping metagenomic reads to redundant genomes within the database, we construct a database of genome representatives for read assignment, similar to [9]. The representative database consists of the first genome from each cluster defined by ANI 99% and is indexed using bowtie2-build [11] with standard parameters. SPARSE indexes 20, 850 bacterial representative genomes in ~ 4 hours using 20 computer processes. Representative databases of other ANI levels or clade-specific databases can also be built by altering the parameters. Furthermore, traditional read mapping tools such as bowtie2 [11] show reduced sensitivity for divergent reads. This is not a problem for many bacterial species, especially bacterial pathogens, because these organisms have been selectively sequenced. However, fewer reference genomes are available for environmental bacteria and eukarya. In order to map reads from such sources to their distantly related references, SPARSE also provides an option to use MALT [5], which is slower than bowtie2 and needs extensive computing memory, but can efficiently align reads onto references with *<*90% similarity.

## 3 Metagenomic read sampling

Given read mappings to the representative databases as input, we adapt a probabilistic model reconstructing the process of sampling reads from a metagenomic sample to assign reads onto reference genomes. We extend the model implemented in Sigma [1] by also considering that reads aligned to a genome in the reference database could still be originating from an unknown source, thus avoiding to overestimate the number of genomes present in the sample. We introduce a weighting for each read reflecting the probability to be sampled from an unknown genome, and show in Section 4 how this improves the precision of taxonomic assignments.

Let *E* denote the set of both known and potentially unknown genomes in a metagenomic sample, and the set of reference genomes included in the SPARSE database is a subset *G ∈ E*. Let *Pr*(*r*_*i*_|*E*) be the probability of sampling a random read *r*_*i*_ from any possible source, we have

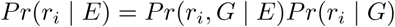

We denote *w*_*i*_ = *Pr*(*r*_*i*_, *G|E*) as the *sampling probability*, indicating the probability that *r*_*i*_ is sampled from any known reference genome in *G*. On the other hand, *Pr*(*r*_*i*_ | *G*) is the probability of generating *r*_*i*_ given *G* and can be further separated as

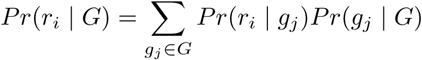

where *Pr*(*g*_*j*_|*G*) is the probability that a genome *g*_*j*_ ∈ *G* was chosen to generate the read, and *Pr*(*r*_*i*_|*g*_*j*_) is the probability of obtaining read *r*_*i*_ from *g*_*j*_. As in Sigma, given a uniform mismatch probability *σ* = 0.05, *Pr*(*r*_*i*_ *g*_*j*_) can be directly calculated from the alignment of *r*_*i*_ to genome *g*_*i*_ with *x* mismatches, and can be stored in a matrix *Q*, such that

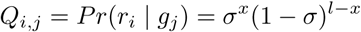

where *l* is the length of read *r*_*i*_. We next describe how the sampling probability *w*_*i*_ is inferred, by giving a weight to each read that indicates the probability of being sampled from a known reference genome. Reads with a low weight do not influence the optimization process used to infer the optimal *Pr*(*g*_*j*_|*G*) for a complete metagenomic read dataset.

### 3.1 SPARSE sampling probability

We model two scenarios that can lead to non-specific mappings of foreign reads.

1) Since there is no systematic way of masking all mobile elements in a reference sequence, we evaluate the probability of a read being drawn from the core genome. We assume that highly conserved regions are part of the core genome, which has been vertically inherited, whereas variable regions likely represent horizontal gene transfers (HGTs). We denote this *HGT probability* as *m*_*i*_.

2) We evaluate the probability of a read originating from an Ultra-Conserved Element (UCE), by comparing the read depths of the aligned genome fragments with other regions in the genome. UCEs are so highly conserved that additional reads from divergent genomes are likely to map on to them, which results in a higher read depth than other regions. We denote this *UCE probability* as *n*_*i*_. Combining both cases as a joint probability, we infer a weight *w*_*i*_ for each read as

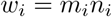

#### HGT probability

Given any cluster *t* in ANI level *k* that consists of *u* references, a read *r*_*i*_ can be assigned to either the core genome *g*_*c*_ or accessory genome *g*_*a*_ of this cluster. Given the number of references *v ⊆ u* the read aligns to, we can formulate the probability of the read originating from the core genome as

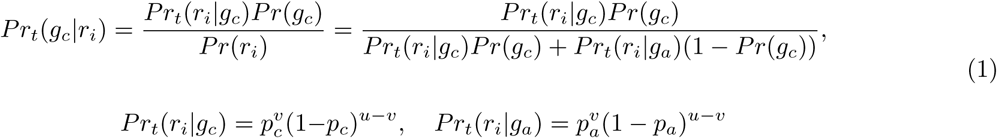

where *Pr*(*g*_*c*_) is the prior probability of any read originating from a core genomic region, and *p*_*c*_ and *p*_*a*_ are the respective probabilities for core genomic fragments or accessory genomic fragments. Default prior probabilities in SPARSE are given in Table 1. Furthermore, a read can align to multiple clusters in the same ANI level *k*, so we average the probabilities of all such clusters for each read weighted by *Q* inferred from the read alignment:

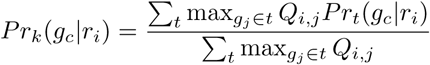

**Table 1:** Default prior probabilities for three ANI levels, values inferred from [3].

| ANI | $Pr(g_c)$ | $p_c$ | $p_a$ |
| --- | --- | --- | --- |
| 90% | 0.05 | 0.99 | 0.1 |
| 95% | 0.02 | 0.99 | 0.2 |
| 98% | 0.01 | 0.99 | 0.5 |

Finally, we consider three different ANI levels for the core genome analysis (by default 90%, 95% and 98%), assigning a lower value for *m*_*i*_ if the read does not map to the core genome at any of these ANI levels:

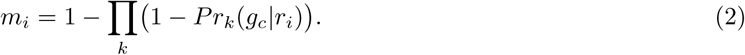

Default values for the prior probabilities were inferred from a published study of core genes across multiple bacterial species [3]. We account for 1% of random deletions of core genes, which gives *p*_*c*_ = 0.99. We also observed that *<*10% of all genes are core genes in bacterial species represented by many genomes. This results in Σ *Pr*(*g*_*c*_) *<* 0.1 over all three ANI levels. We arbitrarily assigned a higher *Pr*(*g*_*c*_) for levels with lower ANI, because a sequence fragment is less likely to be part of a mobile element if it is coincidently present in more divergent genomes. Finally, ~40% of the genes in a random genome are core genes. This gives *m*_*i*_ *≈* 0.6 when *v* = 1 and *u* = 1, which can be used to find empirical values of *p*_*a*_ via equations 1 and 2.

#### UCE probability

In order to compare the read coverage of each fragment in a reference genome *g*_*j*_ with other fragments of the same genome, we split its sequence into *k* consecutive fragments *f*_*j,k*_ using two uniform arbitrary lengths, 487 bps and 2000 bps. Here 487 is used because it is a prime, such that the ends of two fragments overlap only once per Mbp. Then the read depth in each fragment, *d*_*k*_, follows a Poisson distribution with parameter *λ* as the average number of reads per region and probability mass function *f*(*k, λ*). Because of the complexity of the read alignments, we relax the probability of read depth in each fragment such that a wide range of read depths retain high probabilities:

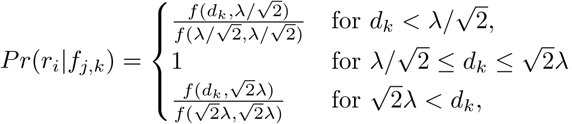

Since a read can again align to multiple genomes *g*_*j*_, we compute the UCE probability of a read as a weighted average of all its alignments. If a read aligns multiple times to the same genome *g*_*j*_ with equal alignment score, we choose one fragment randomly. The UCE probability is then defined as

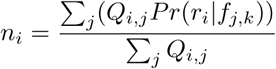

Thus a lower value of *n*_*i*_ is the result from a deviation of the general coverage at the read position in comparison to the average coverage in the genome, indicating that the read is likely mapping to an ultra-conserved region in the genome.

### 3.2 Optimization problem

Knowing the weight *w*_*i*_ for all reads *r*_*i*_ in a whole metagenomic read set *R*, the task is then simplified to finding optimal *Pr*(*g*_*j*_|*G*) values that maximize the probability of the whole read set:

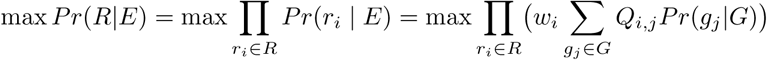

The optimization problem can be solved by a non-linear programing (NLP) method. In SPARSE, we rely on a modified version of the function provided in Sigma [1].

After optimizing *Pr*(*g*_*j*_|*G*), we finally assign a read to a potential reference by checking the following ratio of the computed probabilities:

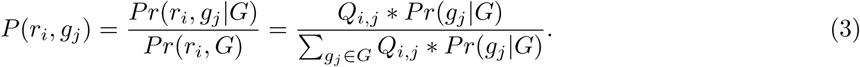

We may assign a read to multiple references, as long as 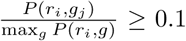. This allows a better abundance estimation for multiple strains from the same species, in which case a read cannot be assigned unambiguously to a single reference.

Further, let *r*_*i*_ ∈ *B* ⊂ *R* be all reads assigned to *g*_*j*_. For a read *r*_*i*_ of length *l* with *x* mismatches in the alignment to its assigned reference, we have a nucleotide similarity of 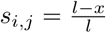. The weighted average similarity 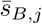 can be calculated as

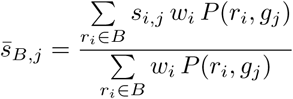

Potentially, reads assigned to a single reference could still originate from several co-existing genomes, with varying degrees of diversity, in the metagenome. We can identify reads from more divergent sources by comparing *s*_*i,j*_ to their average similarity. If all reads assigned to a single reference originate from the same genome in the metagenome, we assume that the similarity of most reads complies with the average similarity over all reads. However, reads originating from very conserved regions show higher similarity than the average and provide a sampling bias. On the other hand, reads originating from different more divergent genomes, will show lower similarity which can be used to avoid overestimating the abundance of each cluster. Therefore we compute the expected average nucleotide identity *s′* for *r*_*i*_ as

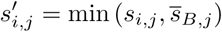

This similarity reflects the ANI between each read and the assigned reference and, as described in the next section, can be used to compute the abundance of each cluster in the metagenomic sample.

### 3.3 ANI cluster abundances

The equation *m*_*i*_*n*_*i*_*P*(*r*_*i*_, *g*_*j*_) describes the probability, for each read *r*_*i*_ ∈ *R*, to be drawn from a region in reference *g*_*j*_ that is part of the core genome (*m*_*i*_) and has even read depth in comparison to the whole chromosome (*n*_*i*_). In summary for all reads assigned to *g*_*j*_, *Σ*_*i*_ *m*_*i*_*n*_*i*_ ∗ *P*(*r*_*i*_, *g*_*j*_) gives the frequency of reads originating from the core genome of *g*_*j*_. However, the desired read abundance for a reference *g*_*j*_ needs to also include reads from the accessory genome. Such reads have been previously suppressed when computing *m*_*i*_. If we assume that all species have the same proportion of core genome, the relative abundances of their core genomes will be equal to the relative abundance of their whole genomes. However, since this is not the case [3], we need to normalize each *m*_*i*_ computed previously. Given *P*(*r*_*i*_, *g*_*j*_) from Equation 3, for any ANI 90% cluster *t*, we normalize *m*_*i*_ for a read *r*_*i*_ as

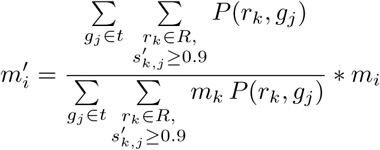

Finally, we assign reads into clusters of all ANI levels according to the references contained in the cluster. For each cluster, we only assign reads if its similarity complies with the ANI level *l* of the cluster, i. e. 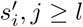.

Thus the abundance of a cluster *t*_*l*_ is computed as the sum of all read abundances assigned to all genomes in the cluster weighted by their probability to originate from an unknown genome. Therefore clusters containing only reads with small *n*_*i*_ and *m*_*i*_ probabilities will receive a low abundance value even if many reads are assigned to it.

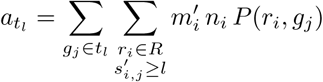

### 3.4 Taxonomic labels for ANI clusters

We finally assign standard taxonomic designations to all clusters at all ANI levels, in order to interpret their biological meaning. Here we rely on a majority vote of all genomes in a cluster. However, the taxonomic levels are restricted to certain ANI levels. For example, species are distinguished at the ANI 95% level, and a species designation is therefore inappropriate for an ANI 90% cluster. Similarly, the taxonomic label for an ANI 95% cluster should not include any subspecies designations.

## 4 Evaluation

### 4.1 Representative Database

We ran SPARSE to index the RefSeq database that consists of 101, 680 complete or draft genomes into 28, 732 clusters at ANI 99% level, which were further grouped into 18, 205 clusters at 95% ANI level, as shown in Fig. 1. Grouping all the genomes according to their species, the resulting representative database is much more evenly distributed, with a Pielou’s evenness [19] of *J′* = 0.9, comparing to *J′* = 0.51 for the whole RefSeq database. Over-representation of pathogenic organisms in the RefSeq database are largely due to repeated sequencing of nearly identical genomes rather than sequencing of intra-species genetic diversities. In particular, nearly half of the genomes in RefSeq are from the top 10 most sequenced bacterial species, which are all human pathogens. All these genomes were grouped into 615 clusters at ANI 99% level, which gives a 65-fold reduction of the data indexed for these species.

**Fig. 1:**
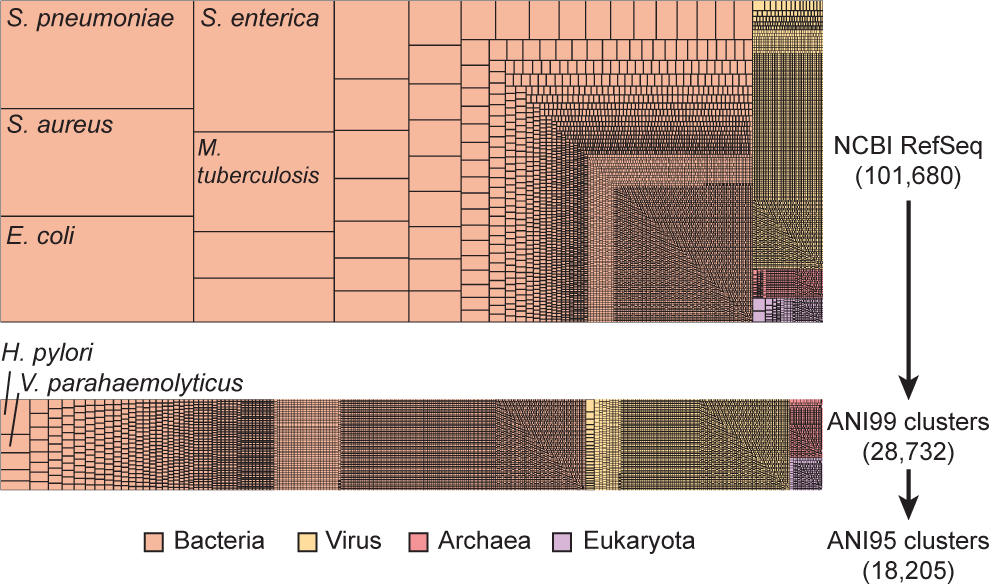
Hierarchical clustering of 101, 680 genomes in NCBI RefSeq database (Aug. 2017) into 18, 205 ANI 95% clusters using SPARSE. Each rectangle represents such a cluster at ANI 95% level, with its area relative to the total number of genomes (top) or clusters at ANI 99% (bottom).

### 4.2 Simulated Data

We ran SPARSE on three recent simulated datasets (Sczyrba et al. [23], McIntyre et al. [14] and Quince et al. [21]). For a fair comparison, the analyses for all datasets were based on a database built from NCBI RefSeq and taxonomy databases dated 22th June, 2015, which is the deadline for the comparison in [23] and also pre-dates the other two comparisons. We evaluated the performance of SPARSE as described in the respective papers for the read-level taxonomic binners, adopting their results for the compared methods. We also included Sigma using the same database as SPARSE in the comparison. We calculated sensitivity and precision based on the number of true-positives (TP; correctly assigned reads), false-positives (FP; incorrectly assigned reads), and false-negatives (FN; unassigned reads).

All simulated reads in the McIntyre et al. [14] study were generated from published complete genomes. This dataset is suitable for comparing the completeness of the databases, as well as the sensitivity of the read mapping approaches in different tools. Both SPARSE and Sigma were run on 18 samples that have read-level taxonomic labels. SPARSE binned all the samples in 10 hours with 20 processes. The precision and sensitivity of both tools in addition to six binning tools from [14] are summarized in Figure 2A. As expected, all tools reached a high precision of > 97%, but differed in their sensitivity. Benefiting from the representative database, SPARSE and Sigma assigned the highest numbers of reads into correct species. The difference between the two methods is due to their different strategies in the modeling, where Sigma assigned all reads to their possible references, whereas SPARSE filtered out unreliable mappings. An independent run of SPARSE using the latest RefSeq database (Aug. 2017) assigned slightly more reads into species, but does not improve precision. This database consists of 20, 850 representative genomes, which is *>* 2 fold the number of representatives (9, 707) in RefSeq 2015.

**Fig. 2:**
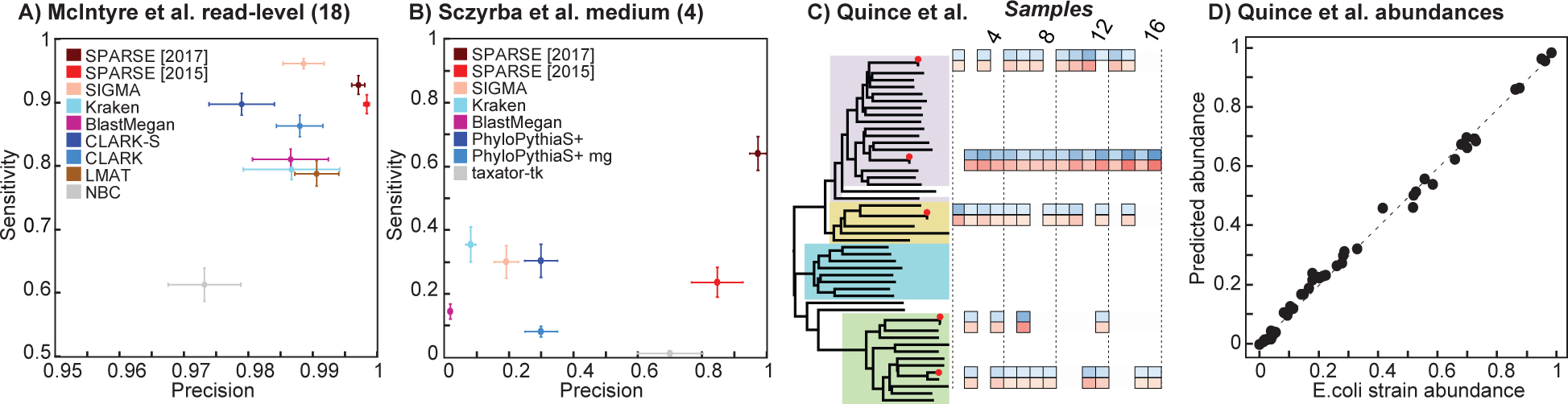
Performances of SPARSE in simulated published datasets. The performance of all the tools in A and B, except for SPARSE and Sigma, are obtained from the respective publications [23, 14]. SPARSE was run in parallel using two different databases. [2015] uses database built from RefSeq at 2015, whereas [2017] uses up-to-date database. A) All the simulated reads in McIntyre et al. [14] were derived from published genomes. B) The Sczyrba et al. [23] used unpublished genomes for read simulations. C+D) Strain-level identification using the mocked *E. coli* datasets as published in [21]. C) Left: The distance-based species tree for*E. coli* for 45 ANI 99% representative genomes plus the five genomes used in [21] for mocked reads. The four largest ANI 98% clusters in *E. coli* are highlighted with colors. Right: Each column shows one of the 16 mocked samples. The true relative abundances of *E. coli* strains in samples (blue) and the relative abundances of predicted strains (red) in samples are shown as colored squares. D) Comparison of true *E. coli* strain abundances versus SPARSE predictions. The dashed line indicates the linear regression of the two values, with *R*^2^ = 0.9948 and *p <* 2.2e−16.

The datasets in Sczyrba et al. [23] are much more challenging, because all the reads were generated from sequencing of environmental isolates, many of which do not have closely related references in the 2015 database. Furthermore, many reads do not have a known microbial species label, because they are not similar to any species in SILVA [20], which was used as the gold standard in this study. We ran both Sigma and SPARSE on the medium complexity datasets, and compared the results with the other methods (see Fig. 2f in [23]) for the recovery of microbial species (Fig. 2B). Using 80 processes, SPARSE ran through all four datasets in ~ 40hours. All the taxonomic binners published in [23] obtained an average precision of *<* 30% at species level, except for taxator-tk [4] with a precision of 70% along with the lowest sensitivity (~ 1.25%). The performance of Sigma is comparable to other binning tools, whereas SPARSE obtained an exceptionally high precision of ~ 85% while still maintaining a sensitivity of ~ 23%. Many incorrect taxonomic bins predicted in Sigma were suppressed in SPARSE, because they have low sampling probability *w*_*i*_ to any of the existing references. Again, SPARSE was also run independently against the database built Aug. 2017. We recovered 63% of the species in the CAMI median datasets, with an average precision of 97%.

Both benchmarks evaluate the performances of taxonomic binnings on or above species level, but give no resolution in intra-species diversity. DESMAN [21] allows reference-free recovery of strain-level variations based on uneven read depths of different strains across multiple samples. It has been compared with two other strain-level binning methods using mock *E. coli* samples [21]. Applying SPARSE to the same 20 genome mocks, we recovered 50*/*51 *E. coli* strains in all 16 samples without any additional strains (false positives), as shown in Fig. 2C. The only strain that was not recovered by SPARSE is 2011C-3493 in the 12*th* sample (Sample733 in [21]), which accounts for only ~ 0.03% of all *E. coli* reads in the sample. We also obtained an almost exact correspondence between the relative abundances of the strains and the predictions (Fig. 2 D). A linear regression of real abundances and the predictions gives an *R*^2^ = 0.9948 and *p <* 2.2e−16.

### 4.3 Ancient Metagenomes

We further evaluated SPARSE and five additional metagenomic tools on three real sets of ancient DNA reads (*Mycobacterium tuberculosis* from [7], *Yersinia pestis* from [22] and *Helicobacter pylori* from [12]) and summarised their results in Table 2. For all samples, the presence of the targeted pathogen, although in very low frequencies (≤ 0.02%), has been confirmed by additional sequencing in the respective publications. MIDAS [16] failed in all three samples and MetaPhlan2 [24] managed to identify *H. pylori* but failed in the other two samples. The results for these two marker-based approaches are consistent with the simulations discussed earlier. Kraken [25] and One Codex [15] are both based on kmer-based taxonomic assignment, but yielded different results. Kraken only identified *H. pylori*, whereas One Codex got positive results in all three samples. However both methods incurred a high number of false positives. For example, Kraken reported *Salmonella enterica* and *Vibrio cholerae* in the Iceman sample, whereas One Codex predicted two *Yersiniae*. All these predictions are inconsistent with results from other tools and analyses presented in the publications. Sigma identified two of three pathogens but inaccurately predicted *V. parahaemolyticus*, which is normally associated with seafood, for the human remains from the Bronze Age. SPARSE successfully identified all three targeted species without any additional suspicious pathogen, which highlights its application to archaeological samples.

**Table 2:**
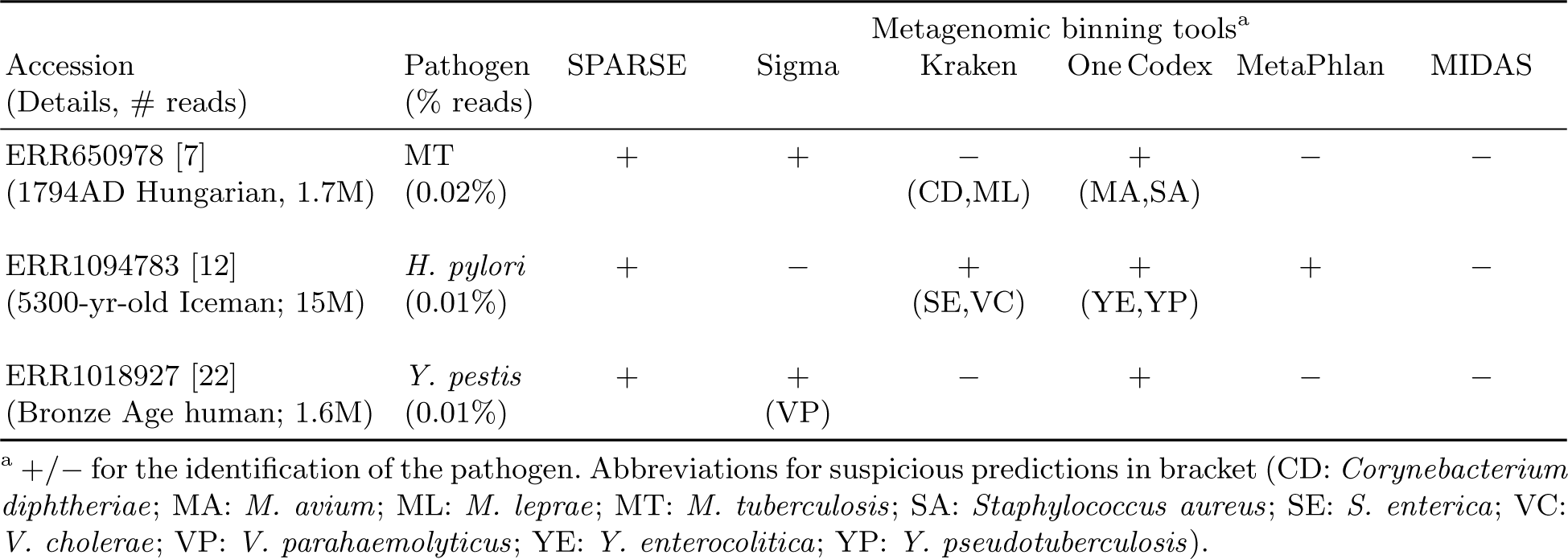
Real archaeological datasets.

## 5 Conclusion

The genetic signatures of specific microbes in metagenomic data, such as human pathogens, are often buried behind the majority of reads from genetically diverse environmental organisms. This is exemplified in the metagenomic sequencing of archaeological samples. Current taxonomic assignment methods compare the metagenomic data with databases that do not fully capture the diversity of microbial genomes. Among these tools, the marker-based taxonomic profilers fail to identify species at low abundances whereas whole genome based taxonomic binners give inaccurate predictions due to non-specific read mappings on ultra-conserved or horizontally transferred elements.

SPARSE indexes existing reference genomes into a comprehensive database with automatic hierarchical clusterings of related organisms. This database is used as a reference for mapping of metagenomic reads. SPARSE penalizes unreliable mappings of reads from unknown sources, and integrates all remaining into a probabilistic model, in which reads were assigned to either an existing reference or unknown sources. In both simulations and real archaeological data, SPARSE outperforms all existing methods, especially in the precisions of species-level assignment. Furthermore, SPARSE managed to identify multiple strains of the same species even when they co-exist in the same sample.

## 6 Acknowledgements

M.A., Z.Z., N.L. and N-F.A. were supported by Wellcome Trust (202792/Z/16/Z). Additional initial grant support was from BBSRC (BB/L020319/1).

## References

1. Ahn, T.H., Chai, J., Pan, C.: Sigma: Strain-level inference of genomes from metagenomic analysis for biosurveillance. Bioinformatics 31(2), 170–177 (2015)

2. Altschul, S.F., Gish, W., Miller, W., Myers, E.W., Lipman, D.J.: Basic local alignment search tool. Journal of Molecular Biology 215(3), 403–410 (1990)

3. Ding, W., Baumdicker, F., Neher, R.A.: panX: pan-genome analysis and exploration. bioRxiv 10.1101/072082 (2016)

4. Dr¨oge, J., Gregor, I., McHardy, A.C.: Taxator-tk: precise taxonomic assignment of metagenomes by fast approximation of evolutionary neighborhoods. Bioinformatics 31(6), 817–824 (2014)

5. Herbig, A., Maixner, F., Bos, K.I., Zink, A., Krause, J., Huson, D.H.: Malt: Fast alignment and analysis of metagenomic dna sequence data applied to the tyrolean iceman. bioRxiv 10.1101/050559 (2016)

6. Huson, D.H., Beier, S., Flade, I., Górska, A., El-Hadidi, M., Mitra, S., Ruscheweyh, H.J., Tappu, R.: MEGAN community edition-interactive exploration and analysis of large-scale microbiome sequencing data. PLoS Computational Biology 12(6), e1004957 (2016)

7. Kay, G.L., Sergeant, M.J., Zhou, Z., Chan, J.Z.M., Millard, A., Quick, J., Szikossy, I., Pap, I., Spigelman, M., Loman, N.J., Achtman, M., Donoghue, H.D., Pallen, M.J.: Eighteenth-century genomes show that mixed infections were common at time of peak tuberculosis in Europe. Nature Communications 6, 6717 (2015)

8. Key, F.M., Posth, C., Krause, J., Herbig, A., Bos, K.I.: Mining metagenomic data sets for ancient DNA: recommended protocols for authentication. Trends in Genetics 33(8), 508–520 (2017)

9. Kim, D., Song, L., Breitwieser, F.P., Salzberg, S.L.: Centrifuge: rapid and sensitive classification of metagenomic sequences. Genome research 26(12), 1721–1729 (2016)

10. Konstantinidis, K.T., Tiedje, J.M.: Genomic insights that advance the species definition for prokaryotes. Proceedings of the National Academy of Sciences 102(7), 2567–72 (2005)

11. Langmead, B., Salzberg, S.L.: Fast gapped-read alignment with Bowtie 2. Nature Methods 9(4), 357–9 (2012)

12. Maixner, F., Krause-Kyora, B., Turaev, D., Herbig, A., Hoopmann, M.R., Hallows, J.L., Kusebauch, U., Vigl, E.E., Malfertheiner, P., Megraud, F., et al.: The 5300-year-old *Helicobacter pylori* genome of the Iceman. Science 351(6269), 162–165 (2016)

13. Marakeby, H., Badr, E., Torkey, H., Song, Y., Leman, S., Monteil, C.L., Heath, L.S., Vinatzer, B.A.: A system to automatically classify and name any individual genome-sequenced organism independently of current biological classification and nomenclature. PLoS ONE 9(2) (2014)

14. McIntyre, A.B.R., Ounit, R., Afshinnekoo, E., Prill, R.J., Hénaff, E., Alexander, N., Minot, S.S., Danko, D., Foox, J., Ahsanuddin, S., et al.: Comprehensive benchmarking and ensemble approaches for metagenomic classifiers. Genome Biology 18(1), 182 (2017)

15. Minot, S.S., Krumm, N., Greenfield, N.B.: One Codex: A Sensitive and Accurate Data Platform for Genomic Microbial Identification. bioRxiv 10.1101/027607 (2015)

16. Nayfach, S., Rodriguez-Mueller, B., Garud, N., Pollard, K.S.: An integrated metagenomics pipeline for strain profiling reveals novel patterns of bacterial transmission and biogeography. Genome Research 26(11), 1612–1625 (2016)

17. O’Leary, N.A., Wright, M.W., Brister, J.R., Ciufo, S., Haddad, D., McVeigh, R., Rajput, B., Robbertse, B., Smith-White, B., Ako-Adjei, D., et al.: Reference sequence (RefSeq) database at NCBI: current status, taxonomic expansion, and functional annotation. Nucleic Acids Research 44(D1), D733–D745 (2015)

18. Ondov, B.D., Treangen, T.J., Melsted, P., Mallonee, A.B., Bergman, N.H., Koren, S., Phillippy, A.M.: Mash: fast genome and metagenome distance estimation using MinHash. Genome Biology 17(1), 132 (2016)

19. Pielou, E.C.: Ecological diversity. Wiley New York (1975)

20. Quast, C., Pruesse, E., Yilmaz, P., Gerken, J., Schweer, T., Yarza, P., Peplies, J., Glöckner, F.O.: The SILVA ribosomal RNA gene database project: improved data processing and web-based tools. Nucleic Acids Research 41(D1), D590–D596 (2012)

21. Quince, C., Delmont, T.O., Raguideau, S., Alneberg, J., Darling, A.E., Collins, G., Eren, A.M.: DESMAN: a new tool for de novo extraction of strains from metagenomes. Genome Biology 18(1), 181 (2017)

22. Rasmussen, S., Allentoft, M.E., Nielsen, K., Orlando, L., Sikora, M., Sj¨ogren, K.G., Pedersen, A.G., Schubert, M., Van Dam, A., Kapel, C.M.O., et al.: Early divergent strains of *Yersinia pestis* in Eurasia 5,000 years ago. Cell 163(3), 571–582 (2015)

23. Sczyrba, A., Hofmann, P., Belmann, P., Koslicki, D., Janssen, S., Dröge, J., Gregor, I., Majda, S., Fiedler, J., Dahms, E., Bremges, A., Fritz, A., Garrido-Oter, R., Jørgensen, T.S., et al.: Critical Assessment of Metagenome Interpretation-a benchmark of metagenomics software. Nature Methods (2017)

24. Truong, D.T., Franzosa, E.A., Tickle, T.L., Scholz, M., Weingart, G., Pasolli, E., Tett, A., Huttenhower, C., Segata, N.: Metaphlan2 for enhanced metagenomic taxonomic profiling. Nature methods 12(10), 902–903 (2015)

25. Wood, D.E., Salzberg, S.L.: Kraken: ultrafast metagenomic sequence classification using exact alignments. Genome Biology 15(3), R46 (2014)

26. Zhou, Z., Lundstrøm, I., Tran-Dien, A., Duchˆene, S., Alikhan, N.F., Sergeant, M.J., Langridge, G., Fotakis, A.K., Nair, S., Stenøien, H.K., et al.: Millennia of genomic stability within the invasive Para C Lineage of *Salmonella enterica*. bioRxiv 10.1101/105759 (2017)

